# Pharmacologic IRE1/XBP1s Activation Promotes Systemic Adaptive Remodeling in Obesity

**DOI:** 10.1101/2020.12.03.408716

**Authors:** Aparajita Madhavan, Bernard P. Kok, Julia M.D. Grandjean, Verena Albert, Ara Sukiasyan, Bibiana Rius, Evan T. Powers, Andrea Galmozzi, Enrique Saez, R. Luke Wiseman

## Abstract

In obesity, overexpression of the IRE1-regulated transcription factor XBP1s protects against metabolic dysfunction by stimulating adaptive remodeling of multiple tissues, most notably the liver.^1–5^ This observation suggests that pharmacologically increasing IRE1/XBP1s signaling might be an attractive approach to mitigate pathologies in obesity and its associated complications.^6–8^ Here, we tested this notion by treating diet-induced obese (DIO) mice with the pharmacologic IRE1/XBP1s activator IXA4.^9^ We show that IXA4 treatment selectively activated protective IRE1/XBP1s signaling in livers of DIO mice without inducing obesity-linked pathologies associated with IRE1 hyperactivity, such as liver inflammation and fibrosis.^10,11^ Chronic IXA4 treatment improved systemic glucose metabolism and feeding-induced insulin action in the liver of DIO mice. These improvements were linked to IRE1/XBP1s-induced remodeling of the liver transcriptome, which dampened glucose production and reduced hepatic steatosis. Further, we show that IXA4 treatment enhanced pancreatic β cell function and insulin homeostasis, indicating that systemic activation of IRE1/XBP1s signaling engendered multi-tissue benefits that integrated to mitigate systemic metabolic dysfunction in DIO mice. Our findings show that selective pharmacological activation of protective IRE1/XBP1s signaling reprograms multiple metabolic tissues, such as liver and pancreas, and represents a potential strategy to correct metabolic alterations in obesity.

## MAIN

The endoplasmic reticulum (ER) transmembrane protein IRE1 regulates the most evolutionarily conserved arm of the unfolded protein response (UPR).^12,13^ In response to ER stress, IRE1 is activated through a mechanism that involves autophosphorylation, oligomerization, and allosteric activation of its cytosolic RNase domain.^12,13^ (**Fig. 1a**). Once activated, IRE1 signals via three mechanisms. IRE1 RNase activity promotes the non-canonical splicing of *XBP1* mRNA that generates the stress-responsive transcription factor XBP1s.^12,13^ IRE1 RNAse activity also promotes promiscuous degradation of mRNA and miRNA through a process called regulated IRE1-dependent decay (or RIDD).^14,15^ Finally, when chronically activated, phosphorylated IRE1 recruits TRAF2 to initiate both JNK-mediated pro-apoptotic signaling and NF-κB-mediated pro-inflammatory signaling.^13,16–18^

**Fig. 1:**
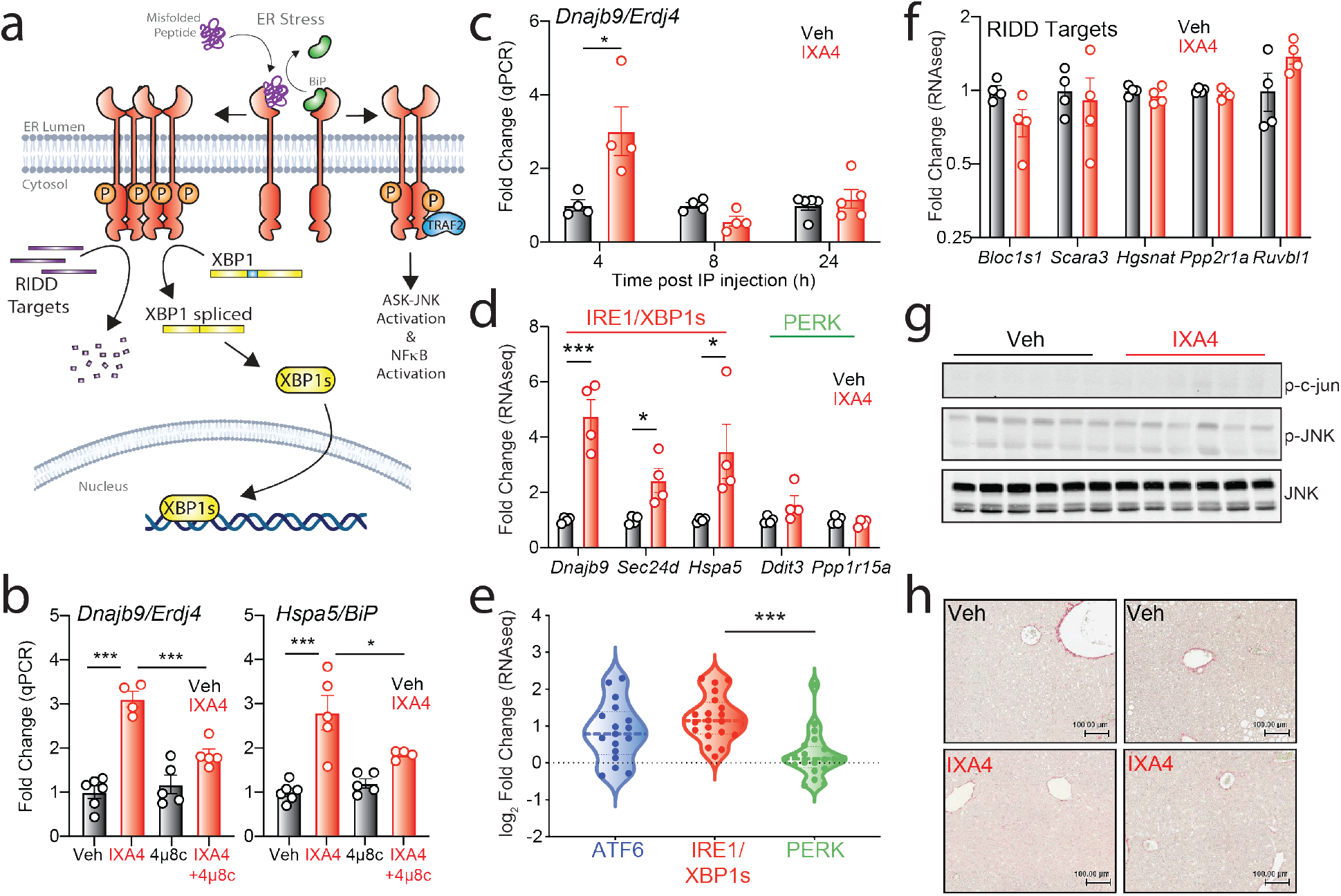
IXA4 selectively induces adaptive IRE1/XBP1s signaling in liver. **a**, Schematic representation of the three downstream consequences of ER stress-induced IRE1 activation: XBP1s-dependent transcriptional signaling, RIDD-associated RNA degradation, and TRAF2-mediated ASK-JNK and NF-κB signaling. **b,** Expression, measured by RT-qPCR, of the XBP1s target genes *Dnajb9* and *Hspa5* in primary mouse hepatocytes treated for 12 h with IXA4 (10 μM) and/or the IRE1 RNase inhibitor 4μ8c (32 μM). **c,** Expression, measured by RT-qPCR, of the IRE1 target gene *Dnajb9* in liver of chow-fed mice 4, 8, or 24 h after acute IXA4 intraperitoneal administration (50 mg/kg). **d,** Fold change in IRE1/XBP1s and PERK target genes, measured by RNA-seq (**Table S1**), in liver of DIO mice after 8 weeks of IXA4 (50 mg/kg) treatment. **e,** Fold change of ATF6, IRE1 and PERK target genesets^30^ in liver of DIO mice after 8 weeks of IXA4 treatment assessed by RNA-seq. Data are shown normalized to vehicle. Specific genes used in this analysis are identified in **Table S2**. **f,** Fold change, measured by RNA-seq, of RIDD targets in liver of DIO mice after 8 weeks of IXA4 treatment. **g,** Immunoblot of c-jun and JNK phosphorylation in liver of DIO mice after 8 weeks of IXA4 treatment. **h,** Representative images of Sirius red staining of DIO mouse livers after 8 weeks of IXA4 treatment. Error bars show SEM for the indicated replicates, *p<0.05, ***p<0.005.

Signaling through these three IRE1-regulated mechanisms is a critical determinant in dictating tissue-specific remodeling in the context of obesity and obesity-linked diseases such as Type 2 diabetes.^1,8,19^ IRE1-dependent XBP1s activation functions primarily to protect tissues such as the liver and the pancreas from the stress brought about by chronic nutrient excess. In obese mice, deletion of *Xbp1* in these tissues causes metabolic dysfunction, inducing hepatic steatosis, insulin resistance, and impairing insulin secretion.^20–24^ In contrast, genetic overexpression of the active XBP1s transcription factor in the liver of obese mice reduces hepatic gluconeogenesis and steatosis and improves systemic glucose metabolism.^2,4,5,25^ Similar improvements in systemic metabolism are observed when *Xbp1s* is overexpressed in other tissues such as adipose depots.^3^ These observations suggest that pharmacologic enhancement of IRE1/XBP1s signaling might constitute a new strategy to mitigate metabolic dysfunction in obesity.

However, prolonged IRE1 hyperactivity is associated with pathologic outcomes in the context of metabolic disease^19^, casting doubt on the viability of this potential pharmacologic strategy. IRE1-dependent JNK signaling promotes inhibitory phosphorylation of insulin receptor substrate (IRS-1) and suppresses insulin signaling in liver cells.^1^ Similarly, hyperactivation of IRE1 in the livers of obese mice is associated with pro-inflammatory signaling and has been implicated in the pathogenesis of hepatocellular carcinoma.^11,26^ Chronic IRE1 hyperactivity also promotes metabolic dysfunction in other tissues such as the pancreas, where sustained IRE1 signaling is associated with both β cell apoptosis and RIDD-dependent degradation of insulin mRNA.^19,27^ Thus, any pharmacologic strategy to target IRE1 signaling in obesity must allow selective increases in protective IRE1 signaling (i.e. *Xbp1* splicing), without inducing pathologies associated with sustained IRE1 hyperactivity (e.g., TRAF2-mediated JNK activation).^6,7,19^

We recently identified a compound, IXA4, that selectively activates protective IRE1/XBP1s signaling in mammalian cells, but does not stimulate RIDD or TRAF2-mediated JNK or NF-κB signaling.^9^ However, the effects of this compound *in vivo* and its potential to improve metabolic function in the context of obesity have not been evaluated. Towards this goal, we initially treated primary mouse hepatocytes with IXA4 and measured expression of XBP1s target genes, including *Dnajb9/Erdj4* and *Hspa5/BiP*. IXA4 treatment induced expression of both these genes (**Fig. 1b**). Importantly, compound-induced increases in the expression of these genes were attenuated by co-treatment with the IRE1 RNase inhibitor 4μ8c^28^, confirming that IXA4 elevated their expression in an IRE1-dependent manner. Next, to test if similar effects would be observed in mice, we dosed chow-fed mice with IXA4 (50 mg/kg; intraperitoneal injection) and monitored expression of the XBP1s target gene *Dnajb9* at multiple time points. IXA4 treatment increased hepatic expression of *Dnajb9* at 4 h, with levels returning to baseline within 8 h of treatment (**Fig. 1c**). These observations indicate that IXA4 treatment induced a transient activation of IRE1/XBP1s signaling in the liver, thus presenting a unique opportunity to test the impact of pharmacologic IRE1/XBP1s activation in the context of obesity.

To evaluate the therapeutic promise of selective, increased IRE1/XBP1s signaling, we administered IXA4 daily to diet-induced obese (DIO) mice for 8 weeks. We then isolated the livers and assessed IRE1/XBP1s activation by RNA-seq and real-time (RT)-qPCR. Similar to what was observed in chow-fed mice, IXA4 treatment induced expression of IRE1/XBP1s target genes (e.g., *Dnajb9)* in DIO mice (**Fig. 1d,e**; **Fig. S1a**, and **Tables S1,S2**). Gene set enrichment analysis (GSEA) confirmed activation of the IRE1/XBP1s transcriptional program in IXA4-treated mice (**Fig. S1b**). IXA4 treatment did not induce genes regulated by the PERK arm of the UPR such as *Chop/Ddit3* or *Gadd34/Ppp1r15a* (**Fig. 1d,e** and **Fig. S1a**). Genes primarily regulated by the ATF6 UPR signaling pathway (e.g., *Hspa5)* were modestly induced by IXA4 treatment; however, the similarity in expression between ATF6 and IRE1/XBP1s target genes reflects the overlap between these transcriptional programs and not ATF6 activation, as previously described.^29,30^ The lack of direct ATF6 activation in livers from IXA4-treated mice was corroborated by the observation that IXA4-dependent induction of *Hspa5*, a gene regulated by both ATF6 and IRE1/XBP1s^29,30^, was blunted by co-treatment with the IRE1 RNase inhibitor 4μ8c in primary mouse hepatocytes (**Fig. 1b**). Collectively, these findings indicate that IXA4 selectively activated the IRE1/XBP1s transcriptional arm of the UPR in the liver of treated DIO mice. Importantly, GSEA confirmed that IXA4 treatment did not activate other stress-responsive transcriptional pathways, such as the heat shock response (HSR) or the oxidative stress response (OSR) (**Fig. S1c,d**). Gene ontology (GO) also indicated that pathways increased by IXA4 treatment were primarily associated with UPR-associated processes including ER stress and secretory proteostasis (**Table S3**), further highlighting the selectivity of this compound in the livers of DIO mice.

Despite the noted increase in IRE1/XBP1s signaling, chronic treatment with IXA4 did not reduce the expression of hepatic RIDD targets (**Fig. 1f**).^31^ Similarly, IXA4 treatment did not increase JNK phosphorylation or downstream induction of pro-apoptotic genes in DIO mice (**Fig. 1g** and **Fig. S1e**). We also did not observe increases in NF-κB transcriptional activity (**Fig. S1f**). These results indicate that IXA4 treatment did not induce RIDD or TRAF2-mediated IRE1 signaling in the liver of treated DIO mice (**Fig. 1a**). This finding likely reflects the transient nature of IXA4-induced IRE1/XBP1s activation in liver (**Fig. 1c**), which is expected to limit the IRE1 hyperactivity associated with these other aspects of IRE1 signaling. Although we noticed an upregulation of select acute phase response (e.g., *Saa1, Saa2* and *Orm2*) and fibrosis genes (e.g., *Col1a1* and *Timp1*) in livers of IXA4-treated DIO mice (**Fig. S1g,h**), IXA4 treatment did not increase plasma cytokines (**Fig. S1i**) or liver fibrosis (**Fig. 1h**). In addition, we observed no increase in mRNA levels of the macrophage marker *Cd68* or of ceramide synthesis genes associated with IRE1-dependent hepatocyte inflammation^32^ in the liver of treated DIO mice (**Fig. S1j,k**). These results indicate that IXA4 treatment did not exacerbate hepatic inflammation or fibrosis. Instead, IXA4 treatment modestly reduced plasma ALT levels, an indication of improved liver function (**Fig. S1l**). Collectively, these findings show that IXA4 selectively stimulated protective IRE1/XBP1s signaling in the liver of DIO mice in the absence of pathologic IRE1 hyperactivity.

Liver overexpression of the active XBP1s transcription factor to physiologically relevant levels improves systemic glucose metabolism in obese-diabetic mice.^2,4,5^ Because IXA4 treatment selectively activated protective IRE1/XBP1s signaling in the liver, we anticipated that this compound might similarly improve systemic glucose metabolism in DIO mice. Indeed, IXA4 treatment reduced fasting blood glucose, plasma insulin, and the HOMA-IR in DIO mice, reflecting improvements in insulin sensitivity (**Fig. 2a-c**). IXA4 treatment also increased systemic glucose clearance measured using a glucose tolerance test (GTT) when glucose was administered either orally (OGTT; **Fig. 2d,e**) or intraperitoneally (IPGTT; **Fig. S2a,b**). Importantly, IXA4-induced enhancement of glucose clearance was independent of changes in body weight or food intake, which were unaffected over the course of treatment (**Fig. S2c,d**). As was observed previously upon hepatic XBP1s overexpression^2^, we noted no increase in insulin-stimulated glucose clearance, as assessed in an insulin tolerance test (ITT), in DIO mice treated with IXA4 for 2 weeks (**Fig. 2f**). AKT phosphorylation was also not affected in gastrocnemius muscle isolated from IXA4-treated DIO mice and acutely challenged *ex vivo* with insulin (**Fig. S2e**). Similarly, AKT phosphorylation was not altered in quadriceps muscle of IXA4-treated DIO mice isolated 15 min after an *in vivo* oral glucose bolus (**Fig. S2f**). However, AKT phosphorylation was increased in livers isolated from IXA4-treated mice after administration of a glucose bolus or a meal challenge (**Fig. 2g,h** and **Fig. S2g**), reflecting improved insulin action in this tissue. These observations indicate that IXA4 improved glucose homeostasis in the liver of treated DIO mice in physiologically relevant contexts such as feeding.

**Fig. 2:**
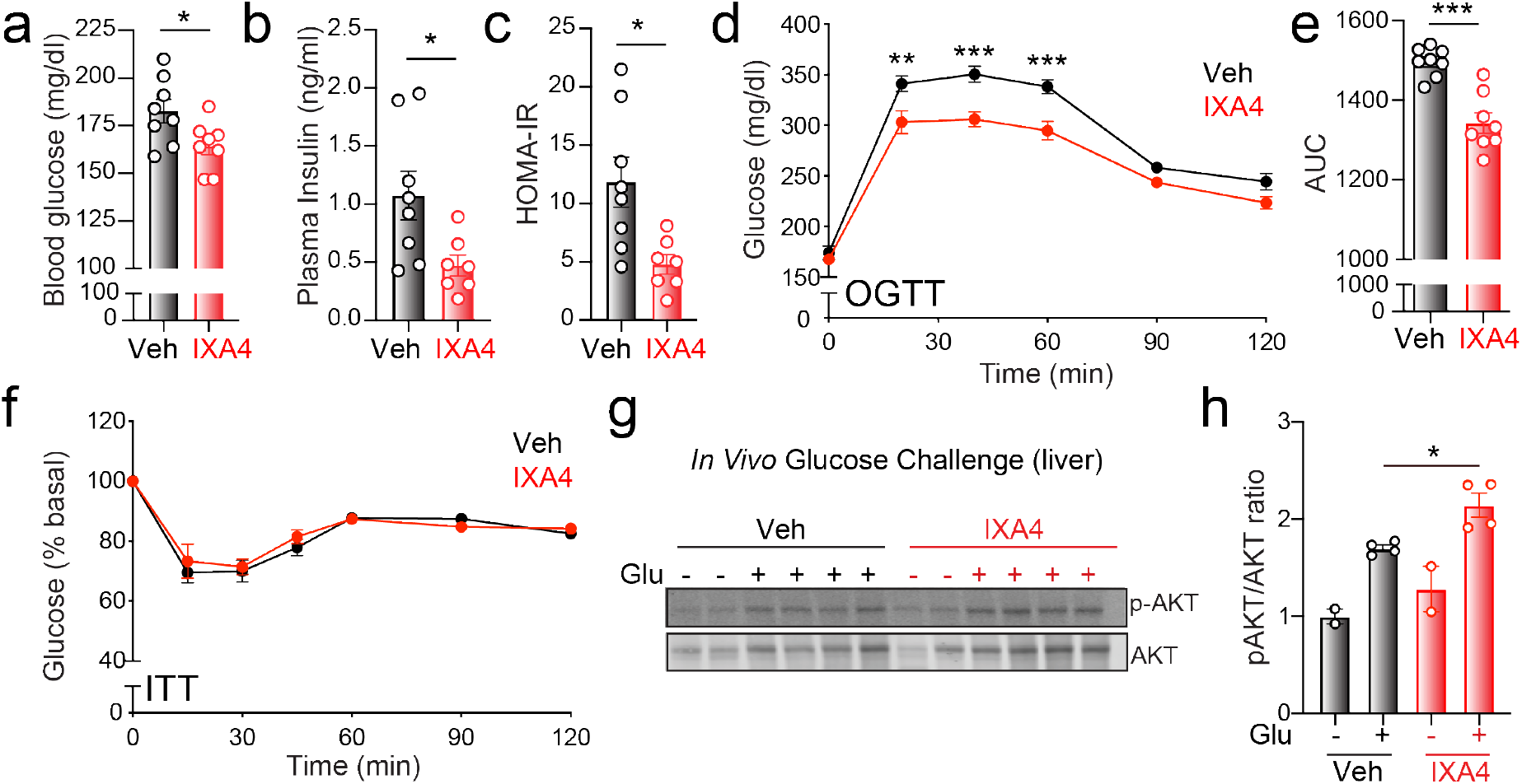
Chronic IXA4 treatment enhances systemic glucose homeostasis. **a-c,** Fasted plasma glucose (**a**), insulin (**b**) and HOMA-IR (**c**) in DIO mice after 46 days of IXA4 treatment. **d,** Oral GTT in DIO mice 38 days after start of IXA4 treatment. Error bars show SEM for n = 8 mice/condition. **e,** Area under the curve quantification of the OGTT in (**d**). **f,** ITT in DIO mice after 17 days of IXA4 treatment. Error bars show SEM for n = 8 mice/condition. **g,** Immunoblot of AKT phosphorylation in liver 15 min after an oral bolus of glucose was administered to DIO mice treated for 60 days with IXA4. **h,** pAKT/AKT ratio of immunoblots shown in (**g**). Error bars show SEM for the indicated replicates, *p<0.05, **p<0.01, ***p<0.005.

Overexpression of XBP1s in the liver enhances hepatic glucose control through two mechanisms: increased expression of hexosamine biosynthesis pathway (HBP) genes^4^, and induction of post-translational degradation of FOXO1 – a key transcriptional regulator of hepatic gluconeogenic gene expression.^2^ These changes act to increase glucose usage through the HBP pathway and to directly suppress hepatic gluconeogenesis by reducing gluconeogenic gene expression. We found that treatment of primary mouse hepatocytes with IXA4 increased expression of HBP target genes (e.g., *Gale, Gfat1*) and reduced FOXO1 protein levels independent of changes to *Foxo1* mRNA (**Fig. 3a**, **Fig. S3a,b**). Both of these IXA4-induced alterations were attenuated by co-treatment with the IRE1 RNase inhibitor 4μ8c, indicating they require IRE1 signaling. Similarly, IXA4 treatment increased expression of HBP genes and reduced FOXO1 protein levels in the liver of chronically treated DIO mice (**Fig. 3b** and **Fig. S3b,c**). However, in this setting, the reduction in FOXO1 protein levels was accompanied by a decrease in *Foxo1* mRNA, likely reflecting the consequences of chronic, long-term treatment with the IRE1/XBP1s activator (**Fig. S3d**). Regardless, and consistent with reduced FOXO1 activity, IXA4 treatment significantly reduced expression of gluconeogenic genes in the liver of DIO mice (**Fig. 3c,d** and **Table S4**). As expected given this observation, pyruvate-dependent hepatic gluconeogenesis, measured using a pyruvate tolerance test (PTT), was reduced in IXA4-treated DIO mice (**Fig. 3e**). These findings show that IXA4 treatment stimulated adaptive remodeling of the liver to reduce hepatic gluconeogenesis.

**Fig. 3:**
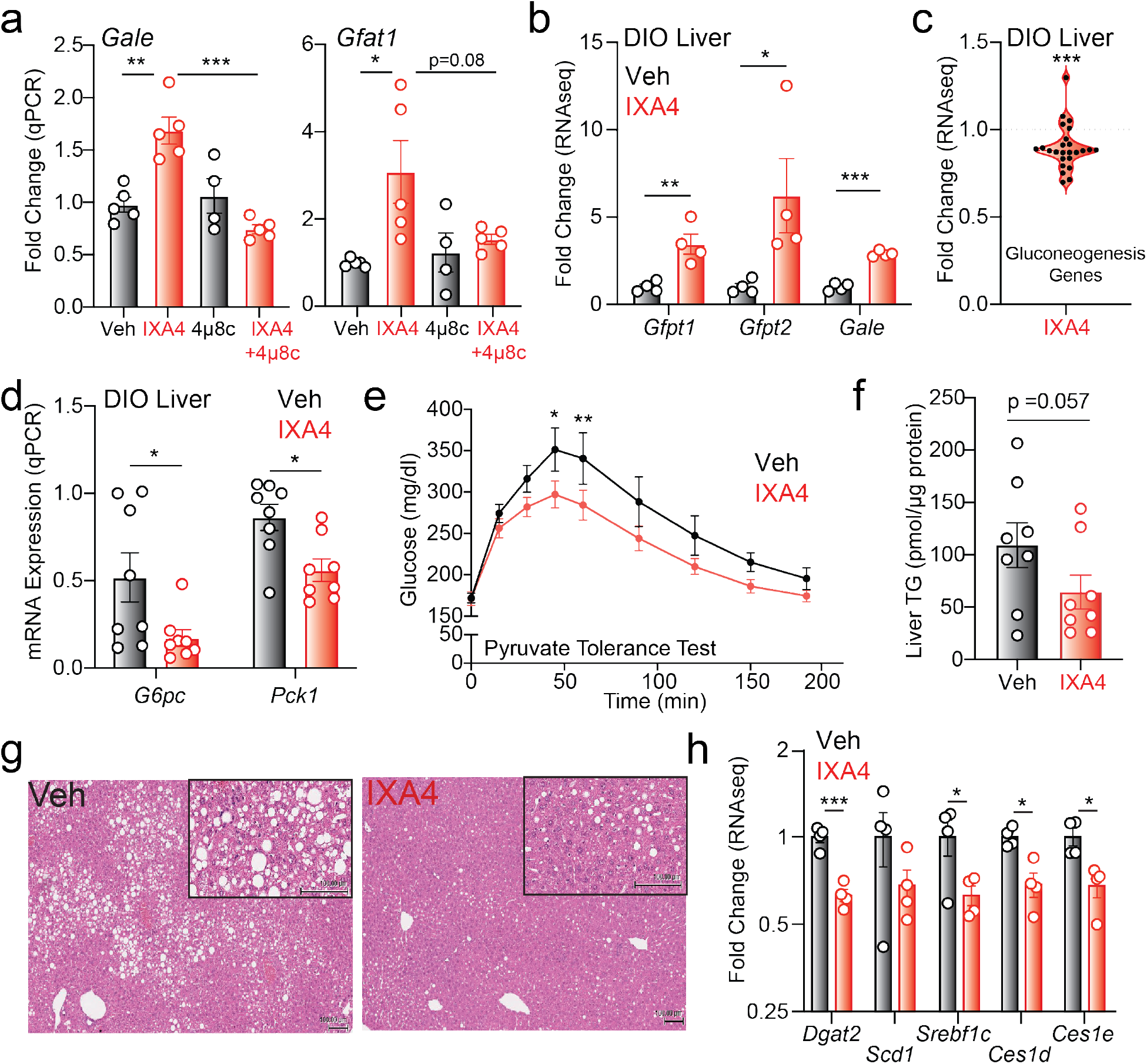
IXA4 improves liver function in DIO mice. **a,** Expression, measured by RT-qPCR, of the HBP genes *Gale* and *Gfat1* in primary mouse hepatocytes treated for 12 h with IXA4 (10 μM) and/or the IRE1 RNase inhibitor 4μ8c (32 μM). **b,** Fold change in HBP genes measured by RNA-seq in liver of DIO mice treated with IXA4 for 8 weeks. **c,** Fold change, assessed with RNA-seq, of gluconeogenic genes in liver of DIO mice treated with IXA4 for 8 weeks. Genes used in this analysis are shown in **Table S4**. **d,** Expression, measured by RT-qPCR, of gluconeogenic genes *G6pc* and *Pck1* in liver of DIO mice treated with IXA4 for 8 weeks. **e,** Pyruvate tolerance test (PTT) in DIO mice treated with IXA4 for 22 days. Error bars show SEM for n = 8 mice/condition. **f,** Triglyceride content in liver of DIO mice treated with IXA4 for 8 weeks. **g,** Representative liver images of DIO mice treated with IXA4 for 8 weeks, stained with H&E. **h,** Fold change, assessed using RNA-seq, of lipogenic genes in liver of DIO mice treated with IXA4 for 8 weeks. Error bars show SEM for the indicated replicates, *p<0.05, **p<0.01, ***p <0.005.

XBP1s also plays an anti-lipogenic role in the liver, suppressing expression of hepatic lipid metabolism genes and reducing liver steatosis.^5^ Accordingly, IXA4 treatment reduced hepatic steatosis and liver triglyceride content in treated DIO mice (**Fig. 3f,g**). These benefits were associated with decreased expression of *Dgat2, Srebp1/Srebf1c*, and *Ces1d* (**Fig. 3h**), all lipid metabolism genes linked to IRE1-dependent suppression of hepatic steatosis.^5,31^ Importantly, the IXA4-stimulated reduction in lipogenic gene expression was blocked in primary mouse hepatocytes co-treated with 4μ8c, indicating that these reductions are cell autonomous and dependent on IRE1 activity (**Fig. S3e**). Thus, in addition to decreasing hepatic glucose production, IXA4 treatment also caused adaptive remodeling of liver lipid metabolism to reduce steatosis in DIO mice.

An advantage of systemic administration of pharmacologic IRE1/XBP1s activators such as IXA4 is the potential to stimulate protective IRE1/XBP1s signaling in multiple tissues. The reductions in fasting plasma glucose and insulin observed in IXA4-treated DIO mice (**Fig. 2a,b**), together with the improved ability of treated mice to clear glucose during a GTT (**Fig. 2d**, **Fig. S2a**), suggested that IXA4 treatment might have also enhanced pancreatic β cell function. Hence, we assessed the effect of IXA4 treatment on the pancreas of IXA4-treated DIO mice. Chronic IRE1 hyperactivation in the pancreas has been shown to alter insulin secretion by promoting islet apoptosis and RIDD-dependent insulin mRNA degradation.^19,27^ However, we observed no change in pancreas morphology or islet number, though we did notice a modest increase in islet size (**Fig. 4a**). IXA4 treatment also did not alter insulin content in isolated islets (**Fig. 4b**). Together, these observations indicate that, as was the case in liver, IXA4 treatment did not increase pathologic IRE1 signaling in the pancreas.

**Fig. 4:**
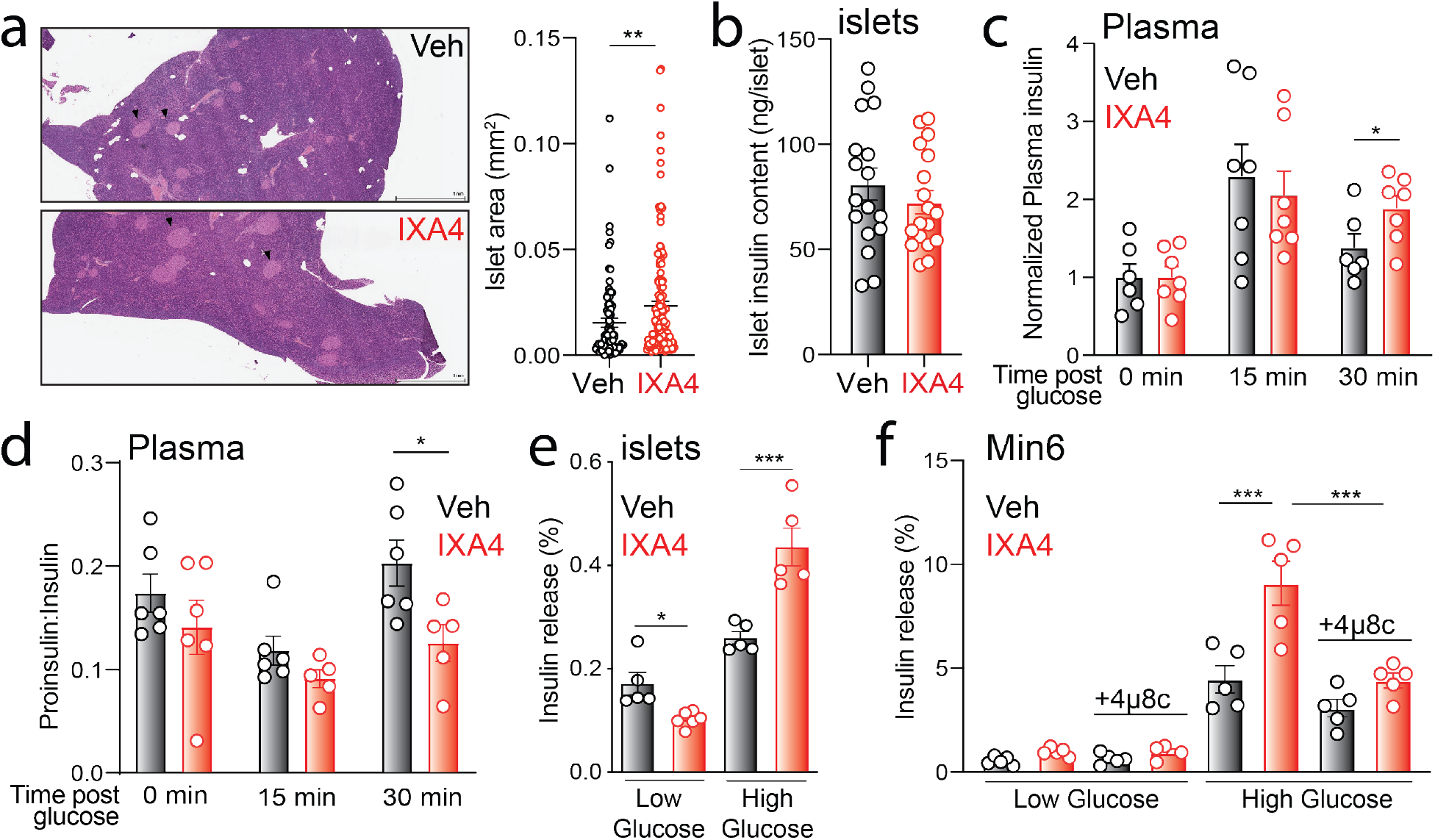
IXA4 treatment enhances pancreatic function. **a,** Representative images and quantification of islet area of pancreas of DIO mice treated with IXA4 for 8 weeks, stained with H&E. Arrowheads indicate individual islets. **b,** Insulin content of primary islets isolated from DIO mice after 8 weeks of IXA4 treatment. **c,** Time course of plasma insulin levels after oral administration of a glucose bolus normalized to basal insulin at time 0 h. Experiment conducted after 38 days of IXA4 treatment. **d,** Plasma proinsulin to insulin ratio after oral administration of a glucose bolus to DIO mice treated for 38 days with IXA4. **e,** Percent insulin release after 60 min of low and high glucose stimulation from primary islets isolated from DIO mice treated for 8 weeks with IXA4. **f,** Insulin release from Min6 cells pretreated for 36 h with vehicle, IXA4 (10 μM), and/or 4μ8c (32 μM) and then stimulated with media containing low (2.8 mM) or high (16.8 mM) glucose for 60 min. Error bars show SEM for the indicated replicates, *p<0.05, **p<0.01, ***p<0.005.

Interestingly, DIO mice chronically treated with IXA4 displayed a more sustained increase in plasma insulin following an oral glucose bolus (**Fig. 4c**). Further, the proinsulin:insulin ratio in plasma was reduced in IXA4-treated mice, indicating greater processing of proinsulin to mature, active insulin following glucose challenge (**Fig. 4d**). These findings pointed to enhanced glucose stimulated insulin secretion (GSIS) in IXA4-treated DIO mice. Consistent with this notion, basal insulin secretion was reduced and GSIS was increased in primary pancreatic islets isolated from IXA4-treated DIO mice (**Fig. 4e**). IXA4-dependent increases in GSIS were also observed in Min6 cells (**Fig. 4f**). Co-treatment with the IRE1 inhibitor 4μ8c, which blocks IRE1-dependent *Xbp1* splicing (**Fig. S4a**), reversed IXA4-induced enhancement of GSIS in Min6 cells, confirming that this effect requires IRE1 activity (**Fig. 4f**). Importantly, we observed no reduction in *Ins1* or *Ins2* mRNA levels in IXA4-treated Min6 cells, indicating that this compound does not induce RIDD-dependent degradation of insulin mRNA in these cells (**Fig. S4b**). Collectively, these results indicate that IXA4 treatment did not spur pancreatic dysfunction associated with chronic IRE1 hyperactivity, but rather, it improved insulin regulation and secretion in the pancreas of DIO mice through adaptive IRE1-dependent reprogramming of pancreatic β cell function.

Our findings reveal the therapeutic potential of pharmacological activation of protective IRE1/XBP1s signaling to stimulate metabolic reprogramming in obesity. We have shown that our small molecule IRE1 activating compound IXA4 induced adaptive IRE1/XBP1s-dependent remodeling in multiple tissues of obese-insulin resistant DIO mice without inducing pathologic IRE1 signaling associated with pathway hyperactivity. Similar to what was observed upon XBP1s overexpression in the liver^2,5^, IXA4 treatment remodeled liver gene expression to reduce hepatic gluconeogenesis and ectopic lipid deposition. Further, we show that IXA4 treatment also improved insulin regulation and secretion in the pancreas, revealing a new mechanism through which pharmacologic IRE1/XBP1s activation aids to restore homeostasis in metabolic disease. These findings provide a compelling rationale for the continued development and application of IRE1/XBP1s activating compounds to mitigate pathologies associated with obesity and other complex diseases.^6,7^

## MATERIALS & METHODS

### Chemicals and Antibodies

The IRE1/XBP1s activator IXA4 was custom synthesized by Otava chemicals. The IRE1 RNase inhibitor 4μ8c was purchased from EMD Millipore (catalog no. 412512). Antibodies used in this study include: phospho-c-jun (Cell Signaling, catalog no. 3270S), phospho-JNK (Cell Signaling, catalog no. 4668S), JNK (Cell Signaling, catalog no. 9252S), phospho-AKT(Ser473) (Cell Signaling, catalog no. 4060S), AKT (Cell Signaling, catalog no. 2920S), FOXO1 (Cell Signaling, catalog no. 2880S), LaminB1 (Cell Signaling, catalog no. 13435S), and tubulin (Sigma, catalog no. T6074-200UL).

### Cell Culture

Min6 cells and primary mouse hepatocytes were cultured in DMEM supplemented with 10% FBS, 2 mM glutamine, 100 U ml^-1^ penicillin and 100 μg ml^-1^ streptomycin in standard culture conditions (37 °C, 5% CO_2_). Primary hepatocytes were isolated from 2-month-old male C57BL/6J mice, as previously described.^33^ Briefly, we cannulated the hepatic portal vein of anesthetized mice and perfused the liver with warm (37°C) 10 mM HEPES-buffered saline for 5 min, followed by 0.05% collagenase in 10 mM HEPES-buffered saline for another 5 min. We then removed the perfused liver, agitated it, and dispersed the cells into culture medium. We filtered the resulting cell suspension through a fine mesh strainer (100 μm) and washed them twice with culture medium. We selected viable cells using Percoll centrifugation, and resuspended them in culture medium before plating them on collagen I-coated plates.

### Mouse Studies

Chow-fed male C57BL/6J mice (10-week-old) were purchased from the Scripps Research breeding colony. Male C57BL/6N diet-induced obese (DIO) mice (14-week-old) were purchased from Taconic Biosciences. DIO mice were maintained on a 60% kcal high fat diet (Research Diets D12492) for 3 weeks prior to IXA4 treatment. IXA4 was formulated in 10% DMSO, 30% Kolliphor EL:Ethanol (2:1 ratio), 60% saline and kept warm until use. Chow-fed or DIO mice were administered IXA4 (50 mg/kg) or vehicle once daily via intraperitoneal injection for up to 8 weeks, as indicated in the text. Body weight and food intake were measured weekly throughout the studies. At sacrifice, tissues were harvested and flash frozen for processing, or collected in formalin for histology. Mouse experiments were approved by and conducted in accordance with the guidelines of The Scripps Research Institute IACUC.

### Glucose, insulin, and pyruvate tolerance tests

For glucose tolerance tests, mice treated with IXA4 or vehicle were fasted overnight for 12 h (21:00 to 09:00) prior to administration of a 2 g/kg glucose bolus via oral gavage or intraperitoneal injection, as noted in the text. For insulin tolerance tests, mice were fasted for 4 h (09:00 to 13:00) prior to intraperitoneal injection of Novolin insulin (0.75 U/kg). For pyruvate tolerance tests, mice were fasted overnight for 12 h (21:00 to 09:00) prior to intraperitoneal injection of sodium pyruvate (1 g/kg). In all cases, plasma glucose levels were determined at the indicated time points using the Clarity BG1000 blood glucose monitoring system (Clarity Diagnostics).

### In vivo insulin signaling upon glucose and meal challenge

To measure insulin signaling upon administration of a glucose bolus or a complex meal, mice were administered 2 g/kg glucose or 200 mL Ensure Plus (catalog no. RS58303), respectively, via oral gavage following a 12 h (21:00 to 09:00) overnight fast. Mice were euthanized 15 min post glucose or meal administration and tissues rapidly excised and frozen for processing and immunoblotting.

### Ex vivo insulin signaling

For measuring insulin signaling upon insulin stimulation *ex vivo*, mice fasted overnight were euthanized and the soleus and gastrocnemius muscles rapidly harvested. Paired tissues were incubated in KHB buffer (138 mM NaCl, 4.7 mM KCl, 1.2 mM MgSO_4_, 1.25 mM CaCl_2_, 1.2 mM KH_2_PO_4_, 25 mM HEPES and 11 mM glucose) supplemented with 0.1% BSA, 2 mM sodium pyruvate, and 6 mM mannitol and containing either vehicle or 10 nM insulin and incubated for 10 min with shaking. Tissues were then washed with PBS and flash frozen in liquid nitrogen until processing and immunoblotting.

### Plasma measurements

Blood glucose levels were determined using the Clarity BG1000 blood glucose monitoring system (Clarity Diagnostics). Blood was obtained by tail vein collection, retro-orbital collection from isoflurane-anesthetized mice, or by cardiac puncture post euthanasia. Plasma insulin was quantified using the Ultra Sensitive Mouse Insulin Elisa (Crystal chem; catalog no. 90080). Plasma proinsulin was measured using the Mercodia Rat/Mouse Proinsulin ELISA (catalog no. 10-1232-01). Plasma cytokines and chemokines were measured using the Bio-Rad Bio-Plex Pro Mouse Cytokine 23-plex Assay (catalog no. M60009RDPD). Triglycerides were quantified using the EnzyChrom Triglyceride Assay (BioAssay systems; catalog no. ETGA-200). Plasma ALT levels were measured using the Amplite Colorimetric Alanine Aminotransferase Assay Kit (AAT Bioquest; catalog no. 13803). HOMA-IR was calculated using the formula: HOMA-IR = glucose (mg/dl) x insulin (mU/L) / 405.

### Primary pancreatic islet isolation and GSIS

Primary mouse islets were isolated from DIO mice treated with IXA4 or vehicle for 8 weeks, as described previously.^34^ Briefly, collagenase P (Roche) was perfused into the pancreas at a concentration of 0.8 mg/ml through the common bile duct. The pancreas was then removed and dissociated at 37°C for a maximum of 15 min. Islets were separated using a gradient composed of HBSS and Histopaque (Sigma) layers. Purified islets were hand-picked using a dissection microscope. Islets from vehicle- or IXA4-treated mice were incubated overnight in RPMI 1640 supplemented with 8 mM glucose, 10% FBS, 2 mM L-glutamine, 100 U/mL Pen/Strep, 1 mM sodium pyruvate, 10 mM HEPES. Next day, islets were washed and pre-incubated for 1 h in Krebs-Ringers-Bicarbonate-HEPES (KRBH) buffer (130 mM NaCl, 5 mM KCl, 1.2 mM CaCl_2_, 1.2 mM MgCl_2_, 1.2 mM KH_2_PO_4_, 20 mM HEPES pH 7.4, 25 mM NaHCO_3_, and 0.1% bovine serum albumin) supplemented with 2.8 mM glucose solution at 37°C in 5% CO_2_. Afterward, groups of 10 islets that were size-matched between groups were transferred to a 96-well plate with KRBH solution containing low glucose (2.8 mM) or high glucose (16.8 mM). After incubation for 1 h, supernatant was collected, and islets were lysed overnight in a 20% acid:80% ethanol solution. Insulin was then measured in supernatants and lysates using the Ultra Sensitive mouse insulin ELISA (Crystal chem). Insulin release was calculated as percentage of total islet insulin content per hour.

### RNA-seq analysis

RNA-seq was performed as previously described.^30^ Briefly, RNA was isolated from livers of DIO mice treated with IXA4 or vehicle (n=4 per condition) using the Zymo Research Quick-RNA Miniprep Kit according to the manufacturer’s instructions. RNA sequencing was performed by BGI Americas on the BGI proprietary platform (DNBseq), providing single-end 50 bp reads at 20 million reads per sample. Alignment of the sequencing data to the *Mus musculus* MGSCv37 NCBI build 37.2 was performed using DNAstar Lasergene SeqManPro. Assembled data were then imported into ArrayStar 12.2 with QSeq (DNAStar Inc) to quantify gene expression and normalized reads per kilobase million (RPKM). Differential expression analysis and statistical significance calculations between conditions were assessed using DESeq in R using a standard negative binomial fit for the aligned counts data and are described relative to the indicated control (**Table S1**). Gene Ontology (GO) enrichment analysis was performed using Panther (geneontology.org) on all transcripts which were significantly (p_adj_ < 0.05) downregulated/upregulated as observed by DESeq. Geneset enrichment analysis (GSEA) was performed using denoted genesets from GO on the GSEA platform for the mouse genome (http://www.informatics.jax.org/vocab/gene_ontology).^35,36^ Genesets used to prepare **Fig. 1e** were defined as previously described and shown in **Table S2**.^30^ Genes used to report on gluconeogenesis in **Fig. 3c** are shown in **Table S4**.

### Real time quantitative PCR (RT-qPCR) analysis

RNA was extracted from cells or tissues using the Zymo Research Quick-RNA Miniprep Kit according to the manufacturer’s instructions. RT-qPCR was performed on complementary DNA synthesized using the High-capacity Reverse Transcription Kit (Applied Biosystems catalog no. 4368814). cDNA was amplified using PowerSYBR Green Master Mix (Applied Biosystems catalog no. 4367659) and primers purchased from IDT. Primer sequences used are listed below:

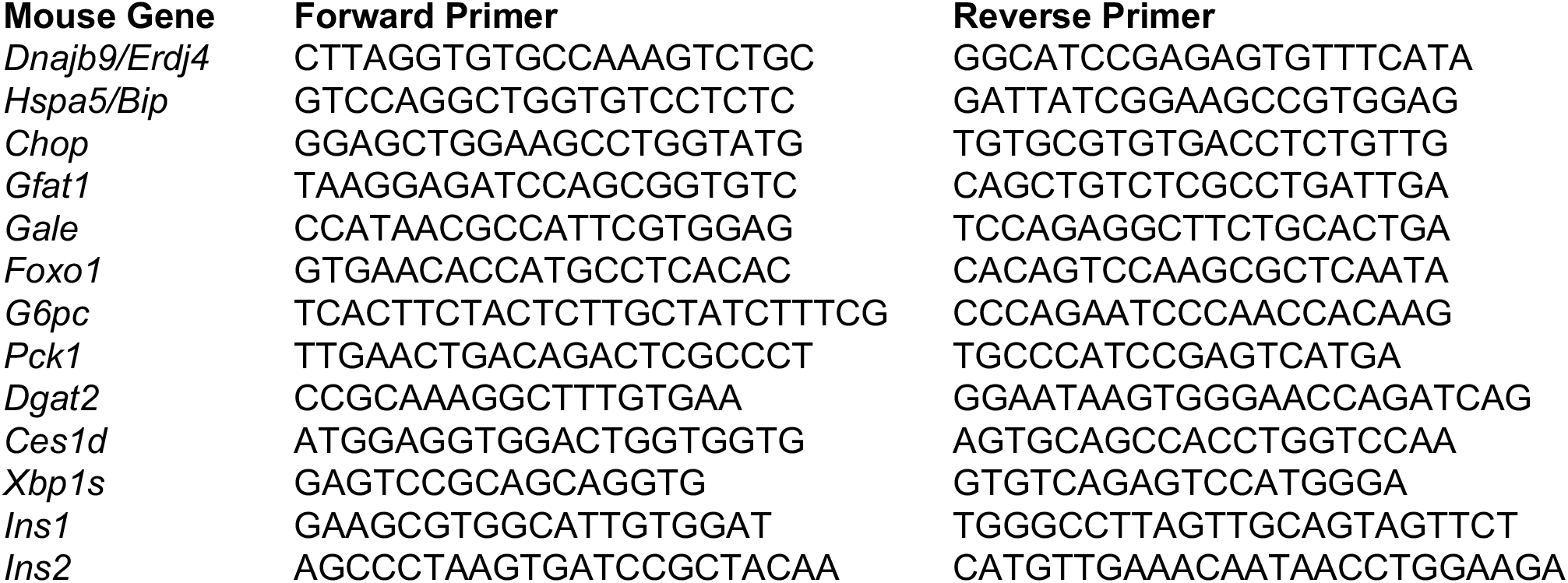

### Immunoblotting

Tissue pieces were homogenized using the Bullet Blender bead homogenizer (Braintree Scientific) and lysed in RIPA buffer (50 mM Tris, pH 7.5, 150 mM NaCl, 0.1% SDS, 1% Triton X100, 0.5% deoxycholate) containing protease and phosphatase inhibitor cocktail (Roche). Total protein concentration was quantified using the Pierce BCA assay kit. Lysates were denatured with 1X Laemmli buffer containing 100 mM DTT and boiled before being separated by SDS-PAGE. Proteins were transferred onto nitrocellulose membranes (Bio-Rad) for immunoblotting and blocked with 5% milk in Tris-buffered saline, 0.5% Tween-20 (TBST) prior to overnight incubation at 4 °C with primary antibodies. Membranes were washed in TBST, incubated with IR-Dye conjugated secondary antibodies and analyzed using the Odyssey Infrared Imaging System (LI-COR Biosciences). Quantification was carried out using LI-COR Image Studio software.

### Histological analysis

Liver and pancreas were processed for histology by the Histology core facility at the Scripps Research Institute. Samples were fixed in formalin, dehydrated, and embedded in paraffin. Tissues were cut into 5 μm thick sections and subject to H&E staining. For Sirius red staining, samples were dewaxed and hydrated prior to incubation with Sirius red solution for 1 h at room temperature. Slides were then washed in acidified water (5 mL glacial acetic acid:1 L distilled water), dehydrated in ethanol and cleared using xylene prior to mounting. Picro-Sirius red solution was purchased from Abcam (catalog no. ab246832). Quantification of islet area from histological sections was performed using ImageJ software.^37^

### Statistics

All data is expressed as mean ± SEM for the indicated replicates. Statistical analysis was performed using a two-tailed Student’s t-test for two groups and one-way ANOVA for more than two groups. A probability value of p < 0.05 was considered significant. Samples were excluded based on outlier testing, as appropriate.

## Acknowledgements

This work was funded by NIH grants AG046495 and DK123038 to RLW, and DK114785 to ES. We thank Jonathan Lin (Stanford) for critical reading of the manuscript. We would also like to thank Jeff Kelly and Kyunga Lee (Scripps) for helpful discussions related to this work.

## Competing Interests Statement

RLW is an inventor on a patent describing IRE1/XBP1s activating compounds, including IXA4. The authors declare no other competing interests.

## Author Contributions

AM, BPK, ES, and RLW designed the research. AM, BPK, JMDG, AG, AS, and BR performed *in vivo* experiments. AM, VA, and BPK performed *in vitro* experiments. AM, JMDG, ETP, and RLW analyzed RNA-seq data. ES and RLW supervised all aspects of this work. AM, BPK, ES, and RLW wrote the manuscript with input from other co-authors.

## Data Availability

The raw data that supports findings within this paper are available from the corresponding authors upon reasonable request. RNA-seq data is available at the public National Center for Biotechnology Information Gene Expression Omnibus repository under the data identifier GSE162567.

## Code Availability

Code for standard open-source DESEQ differential gene expression RNA-seq analysis used in R statistical software is available from the corresponding author upon reasonable request.

---

**Fig. S1:**
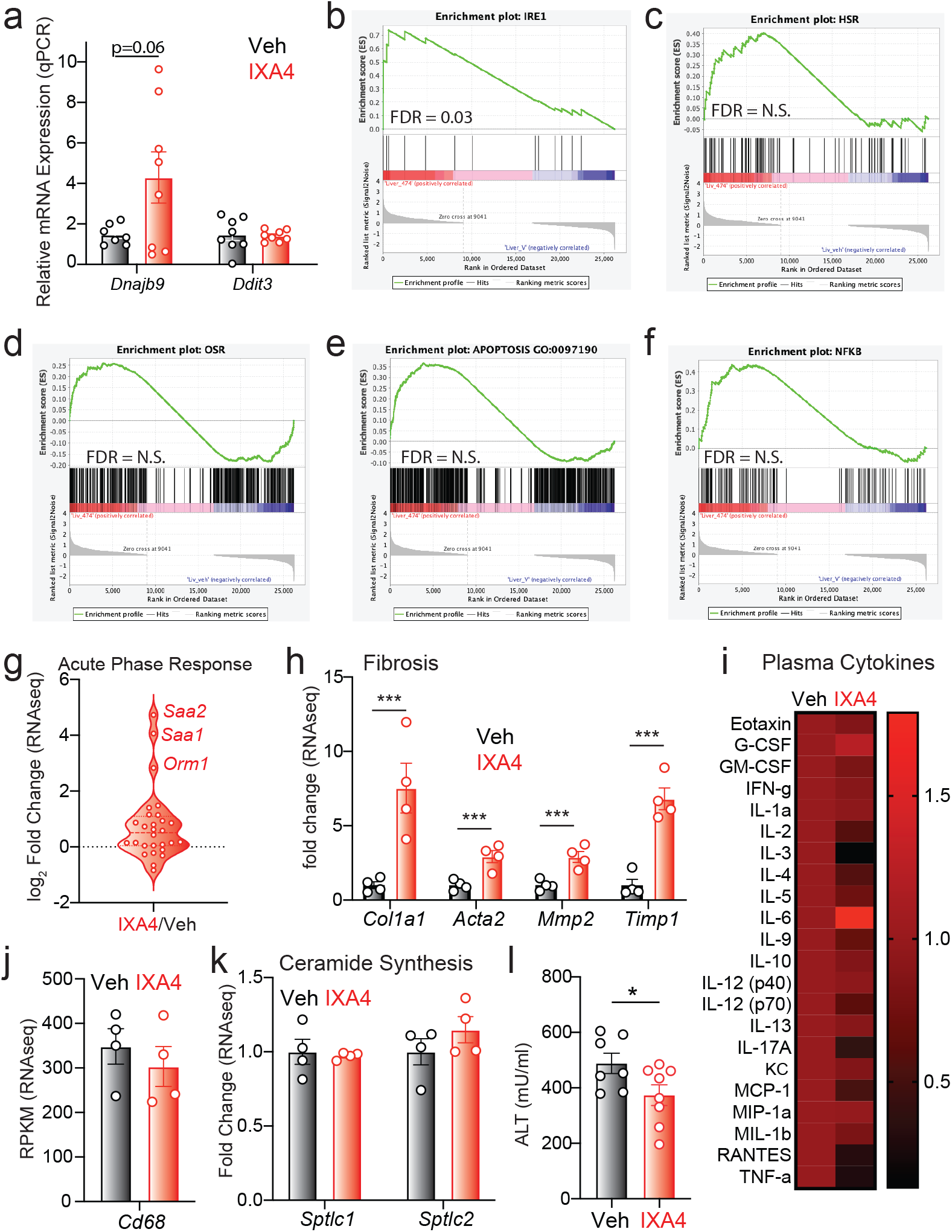
IXA4 is a selective activator of protective IRE1/XBP1s signaling in liver. **a,** Expression, measured by RT-qPCR, of the IRE1 target gene *Dnajb9* and the PERK target gene *Ddit3* in liver of DIO mice treated with IXA4 for 8 weeks. **b-f,** GSEA analysis performed on RNA-seq data from livers of DIO mice treated with IXA4 for 8 weeks for IRE1 signaling (**b**), the heat shock response (**c**), the oxidative stress response (**d**), apoptosis (**e**), and NF-κB signaling (**f**). **g-h,** Fold change, assessed with RNA-seq, of acute phase response geneset (**g**) and fibrosis genes (**h**) in livers of DIO mice treated with IXA4 for 8 weeks. **i,** Heat map of average levels of plasma cytokines (n = 5-19 mice/condition; mice where cytokines fell below the range of detection were excluded from this averaging) from IXA4-treated DIO mice normalized to vehicle. **j,k,** Expression, assessed with RNA-seq, of *Cd68* (**j**) and ceramide synthesis genes (**k**) in liver of DIO mice treated with IXA4 for 8 weeks. **l,** Plasma alanine transaminase (ALT) levels in DIO mice treated with IXA4 for 8 weeks. Error bars show SEM for the indicated replicates, *p<0.05, ***p< 0.005.

**Fig. S2:**
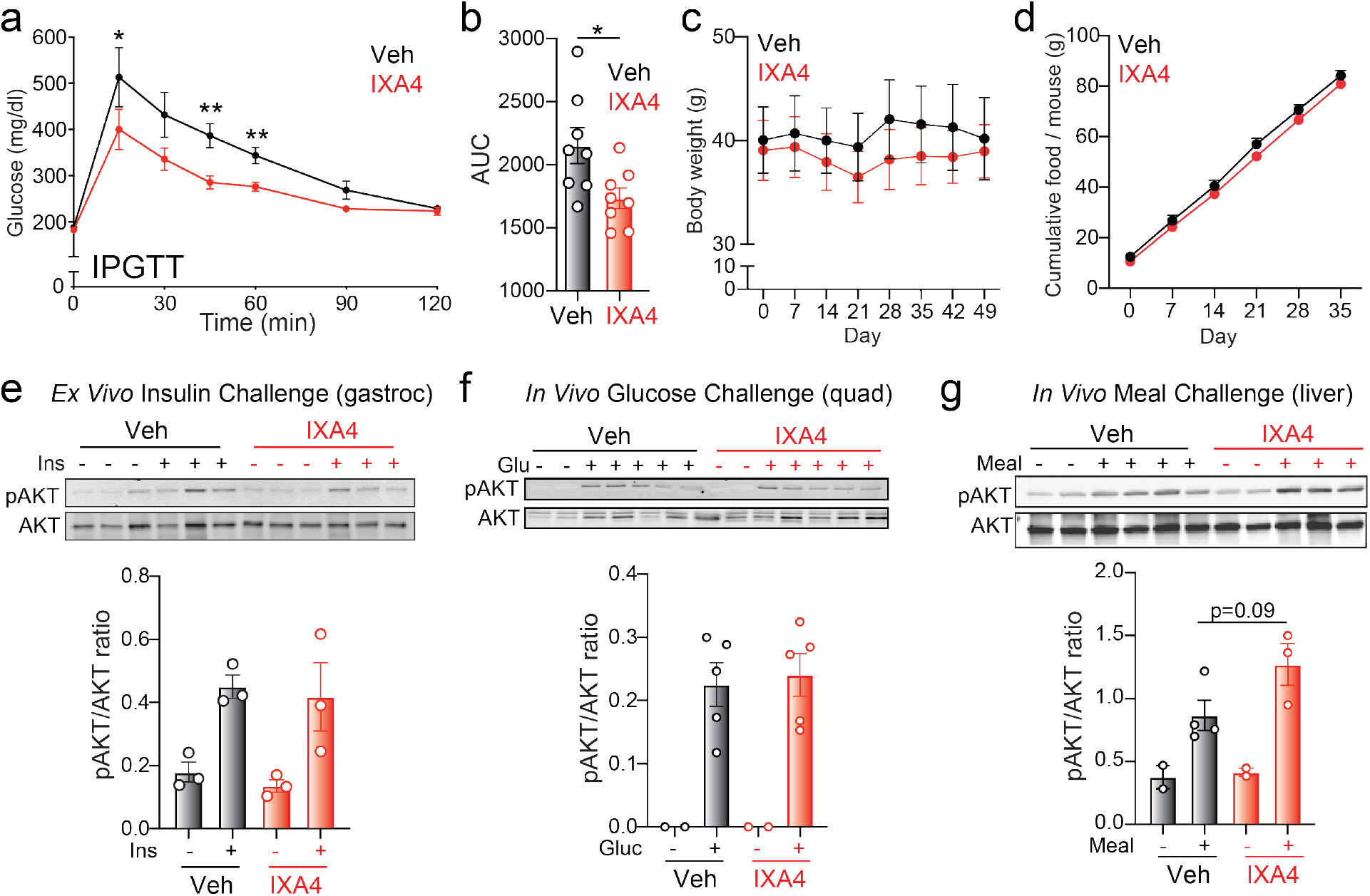
Effects of IXA4 treatment on weight, food intake, and insulin signaling in muscle and liver. **a,** Intraperitoneal GTT (IPGTT) in DIO mice following 22 days of IXA4 treatment. **b,** AUC quantification of (**a**). **c-d,** Body weight (**c**) and food intake (**d**) over time for vehicle and IXA4-treated DIO mice. **e,** Immunoblot and quantification of pAKT/AKT ratio in gastrocnemius muscle isolated from IXA4 or vehicle treated DIO mice and then stimulated for 10 min *ex vivo* with insulin (10 nM). **f,** Immunoblot and quantification of pAKT/AKT ratio in quadricep muscle of DIO mice treated for 60 days with IXA4 or vehicle, 15 min after oral glucose administration. **g,** Immunoblot and quantification of pAKT/AKT ratio in liver of DIO mice treated for 62 days with IXA4 or vehicle, 15 min after administration of a complex meal (Ensure). Error bars show SEM for the indicated replicates, *p < 0.05, **p<0.01, ***p<0.005.

**Fig. S3:**
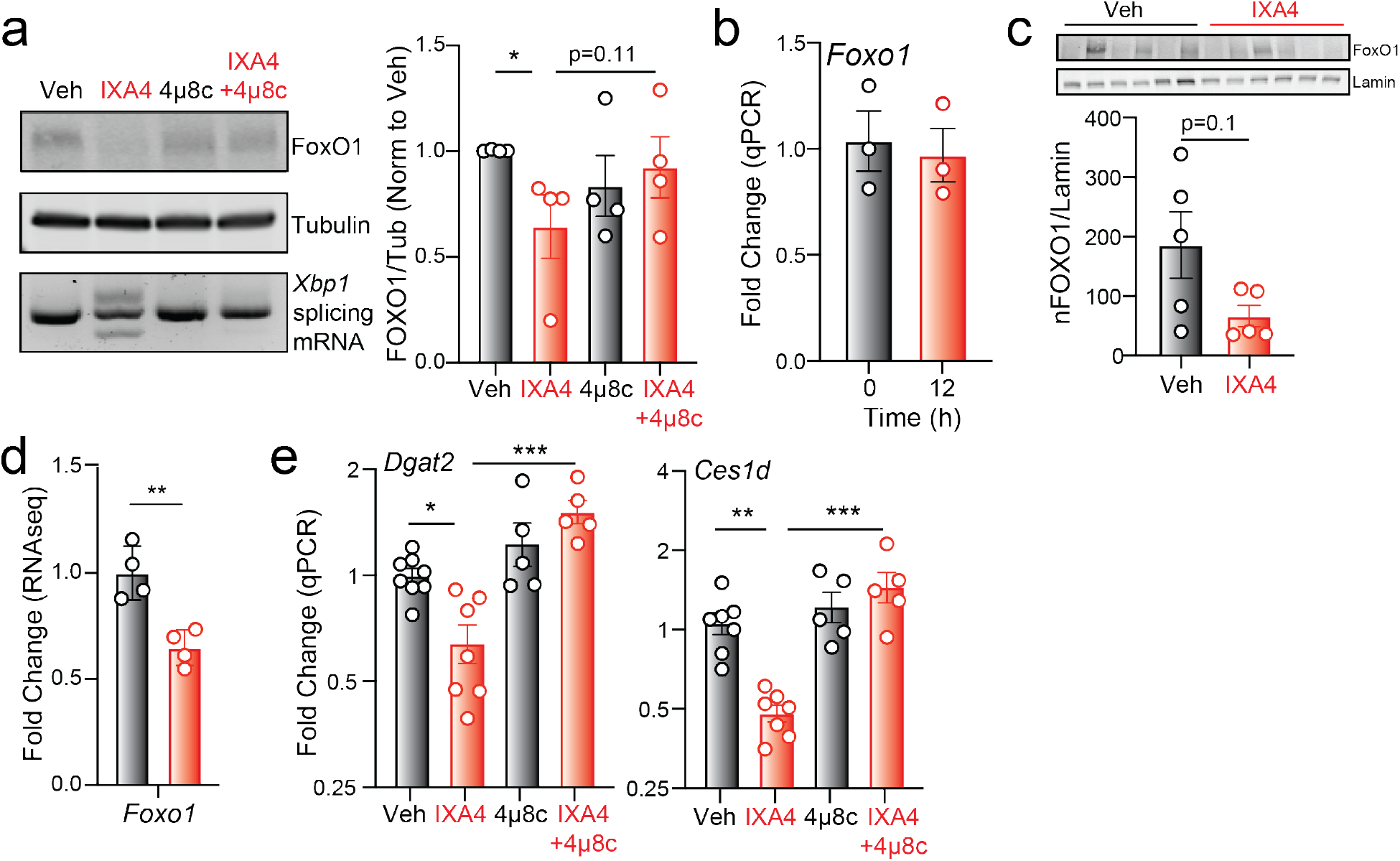
Impact of IXA4 treatment on liver gluconeogenesis and lipogenesis in DIO mice. **a,** Immunoblot and quantification of total FOXO1 protein and tubulin in primary mouse hepatocytes treated for 12 h with IXA4 (10 μM) and/or the IRE1 RNase inhibitor 4μ8c (32 μM). Levels of *Xbp1* splicing are shown in the bottom panel. **b,** *Foxo1* mRNA levels after 12 h of treatment with IXA4 (10 μM) and/or 4μ8c (32 μM). **c,** Immunoblot and quantification of FOXO1 protein in liver nuclear fractions from vehicle and IXA4-treated DIO mice after 8 weeks. **d,** Fold change in *Foxo1* mRNA measured with RNA-seq in liver of DIO mice treated with IXA4 for 8 weeks. **e,** Expression, measured by RT-qPCR, of the lipid metabolism genes *Dgat2* and *Ces1d* in primary hepatocytes after 12 h of treatment with IXA4 (10 μM) and/or 4μ8c (32 μM). Error bars show SEM for the indicated replicates, *p < 0.05, **p<0.01, ***p< 0.005.

**Fig. S4:**
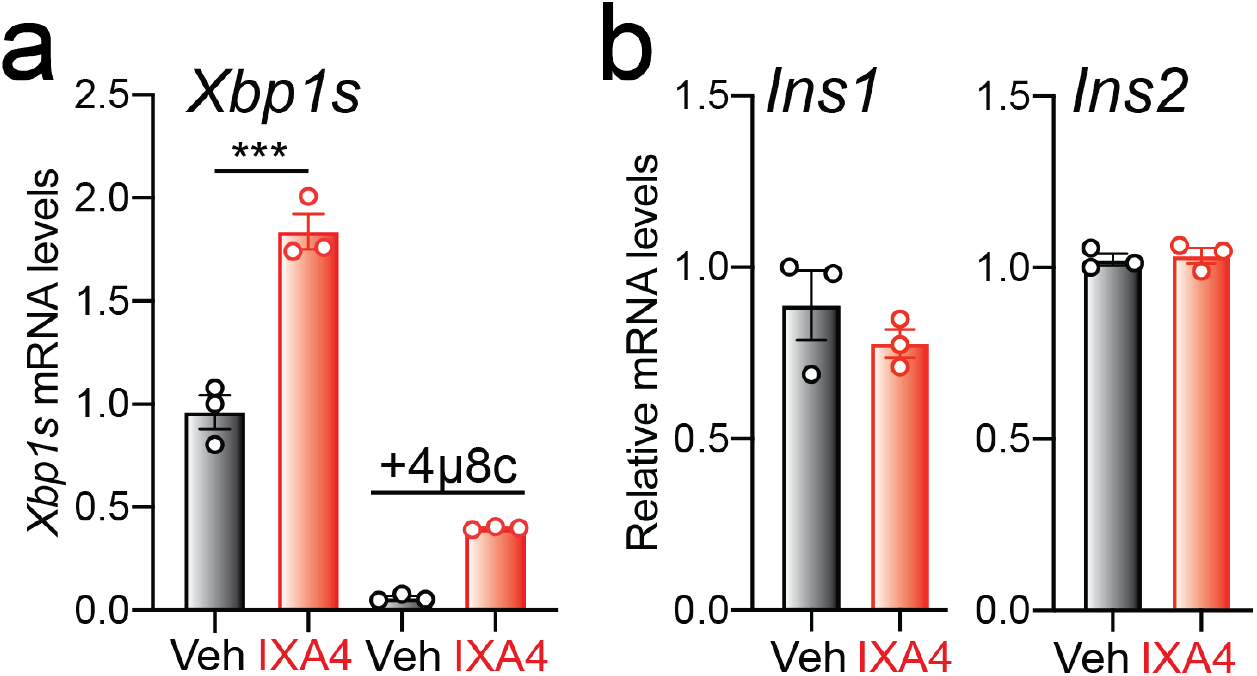
IXA4 activates IRE1/XBP1s signaling in Min6 cells. **a,b,** Expression levels of *Xbp1s* (**a**), *Ins1*, and *Ins2* (**b**) mRNA in Min6 cells after a 6 h treatment with IXA4 (10 μM) and/or the IRE1 RNase inhibitor 4μ8c (32 μM), measured by RT-qPCR. Error bars show SEM for n = 3 replicates, ***p<0.005.

## SUPPLEMENTARY TABLE LEGENDS

**Table S1.** Differential expression (DESeq) analysis of RNA-seq data from liver of DIO mice treated with IXA4 relative to vehicle-treated mice.

**Table S2.** Expression of UPR target genes (measured using RNA-seq) primarily regulated downstream of ATF6, IRE1/XBP1s, or PERK signaling.

**Table S3**. Gene ontology analysis of differentially expressed transcripts (RNA-seq) from livers of DIO mice treated with IXA4 relative to vehicle-treated mice.

**Table S4.** Fold change expression of gluconeogenesis genes (RNA-seq) in livers of DIO mice treated with IXA4 relative to vehicle-treated mice.

